# Imperfect Bayesian inference in visual perception

**DOI:** 10.1101/402776

**Authors:** Elina Stengård, Ronald van den Berg

**Affiliations:** Department of Psychology, University of Uppsala, Uppsala, Sweden

**Author notes:** Corresponding author: (RVDB).

## Abstract

Optimal Bayesian models have been highly successful in describing human performance on perceptual decision-making tasks, such as cue combination and visual search. However, recent studies have argued that these models are often overly flexible and therefore lack explanatory power. Moreover, there are indications that neural computation is inherently imprecise, which makes it implausible that humans would perform optimally on any non-trivial task. Here, we reconsider human performance on a visual search task by using an approach that constrains model flexibility and tests for computational imperfections. Subjects performed a target detection task in which targets and distractors were tilted ellipses with orientations drawn from Gaussian distributions with different means. We varied the amount of overlap between these distributions to create multiple levels of external uncertainty. We also varied the level of sensory noise, by testing subjects under both short and unlimited display times. On average, empirical performance – measured as *d*’ – fell 18.1% short of optimal performance. We found no evidence that the magnitude of this suboptimality was affected by the level of internal or external uncertainty. The data were well accounted for by a Bayesian model with imperfections in its computations. This “imperfect Bayesian” model convincingly outperformed the “flawless Bayesian” model as well as all ten heuristic models that we tested. These results suggest that perception is founded on Bayesian principles, but with suboptimalities in the implementation of these principles. The view of perception as imperfect Bayesian inference can provide a middle ground between traditional Bayesian and anti-Bayesian views.

**Author summary:** The main task of perceptual systems is to make truthful inferences about the environment. The sensory input to these systems is often astonishingly imprecise, which makes human perception prone to error. Nevertheless, numerous studies have reported that humans often perform as accurately as is possible given these sensory imprecisions. This suggests that the brain makes optimal use of the sensory input and computes without error. The validity of this claim has recently been questioned for two reasons. First, it has been argued that a lot of the evidence for optimality comes from studies that used overly flexible models. Second, optimality in human perception is implausible due to limitations inherent to neural systems. In this study, we reconsider optimality in a standard visual perception task by devising a research method that addresses both concerns. In contrast to previous studies, we find clear indications of suboptimalities. Our data are best explained by a model that is based on the optimal decision strategy, but with imperfections in its execution.

## INTRODUCTION

An important function of the visual system is to make inferences about the environment from noisy sensory input. It is often claimed that human performance on perceptual inference tasks is optimal or “Bayesian” [1–5], meaning that subjects supposedly perform as well as theoretically possible given the amount of sensory noise in their observations. Evidence for this claim has mainly come from tasks in which subjects integrate two sensory cues to estimate a common source. The optimal strategy in these tasks is to compute a weighted average of the two cues, where each weight depends on the cue’s reliability: the more reliable the cue, the more strongly it weighs in on the decision [6]. Reliability-based weighting is a hallmark of Bayesian observers and predicts that a subject’s estimates are biased towards the more reliable cue. This prediction has been confirmed in a wide range of experiments in which two sensory cues need to be combined to estimate a common source. Examples include integration of a visual and haptic cue to estimate the height of an object [7], a visual and proprioceptive [8] or auditory [9] cue to estimate object location, and two visual cues to estimate object depth [10,11] or object slant [12]. More recent work has reported that optimality in perception extends to tasks with as many as eight cues and with highly non-linear optimal decision rules, including visual search [13–17], categorization [18], change detection [19], change localization [20], and sameness discrimination [21] tasks.

While these studies have provided valuable insights into basic mechanisms of perception, they have also been criticized. One criticism is that the emphasis on optimality has led to an underreporting and underemphasizing of studies that have found violations of optimality [22,23]. Another, more fundamental criticism is that optimal models often lack explanatory power due to being overly flexible [23–26]. The risk of too much flexibility is that it may allow an optimal model to account for data from suboptimal observers. For example, when sensory noise levels are fitted as free parameters – as in most studies – an optimal model may account for suboptimalities in inference by overestimating these noise levels. Similarly, a freely fitted lapse rate may help an optimal model to explain away errors that were in reality caused by poor decision making. In addition to this methodological concern, several recent studies have suggested that neural computation is inherently imprecise [27–31], which makes it a priori implausible that humans perform optimally on any non-trivial task.

Here, we revisit optimality in perception by using a method that takes note of the concerns described above in three ways. First, we constrain flexibility of the optimal model by imposing prior distributions on its parameters; this reduces the risk that the optimal model explains away decision suboptimalities as sensory noise or attentional lapses. Second, for each model that we test, we also include a variant with computational imperfections. Such imperfections may produce suboptimal behavior, even when subjects use a decision strategy that is based on Bayesian principles. By including these models in our analyses, we can distinguish performance loss caused by using a wrong decision rule form performance loss due to imperfect execution of a rule. Third, besides only testing which kind of model accounts best for behavior, we will also quantify performance loss and partition this loss into different sources (see [27] for a similar approach).

We choose visual search as our experimental task. Despite the complexity of the optimal decision rule for this task, several previous studies have reported that humans perform near-optimally [13–15]. We include experimental conditions in which stimuli are corrupted by external noise, which makes the task more consistent with naturalistic conditions, where inference often involves dealing with both internal and external sources of uncertainty [32,33]. We fit several Bayesian model variants as well as ten heuristic models to the experimental data. To preview our main result, we find no evidence for perfect optimality, nor for any of the heuristic-based strategies. Instead, the data are best explained by an “imperfect Bayesian”, in which decisions are based on Bayesian principles, but subject to imprecisions in the implementation of these principles.

## METHODS

### Data and code sharing

The experimental data are available at https://osf.io/dkavj/. Matlab code that reproduces the main results will be made available upon publication.

### Subjects

Thirty subjects were recruited via advertisements at the psychology department of Uppsala University in Sweden and received payment in the form of cinema tickets or gift vouchers. All subjects had self-reported normal or corrected-to-normal vision and gave informed consent before the start of the experiment. No subjects were excluded from any of the analyses.

### Stimuli

Stimuli were black ellipses (0.35 cd/m^2^) with an area of 0.60 deg^2^ presented on a gray background (71 cd/m^2^; Fig. 1A). The task-relevant feature in all experiments was ellipse orientation, with 0° defined as vertical. The eccentricity of the ellipses differed across stimuli and conditions. Ellipse eccentricity is formally defined as 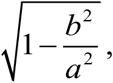, where *a* and *b* specify the ellipse’s semi-major axis and semi-minor axis, respectively. To avoid confusion with visual field eccentricity, we will refer to this eccentricity as “elongation”. Differences in elongation were used to create differences in the level of sensory noise across stimuli (Fig. 1B). Stimuli were generated using the Psychophysics Toolbox [34] for Matlab and presented at fixed locations along an invisible circle at the center of the screen and with a radius of 7 degrees of visual angle.

**Fig. 1.**
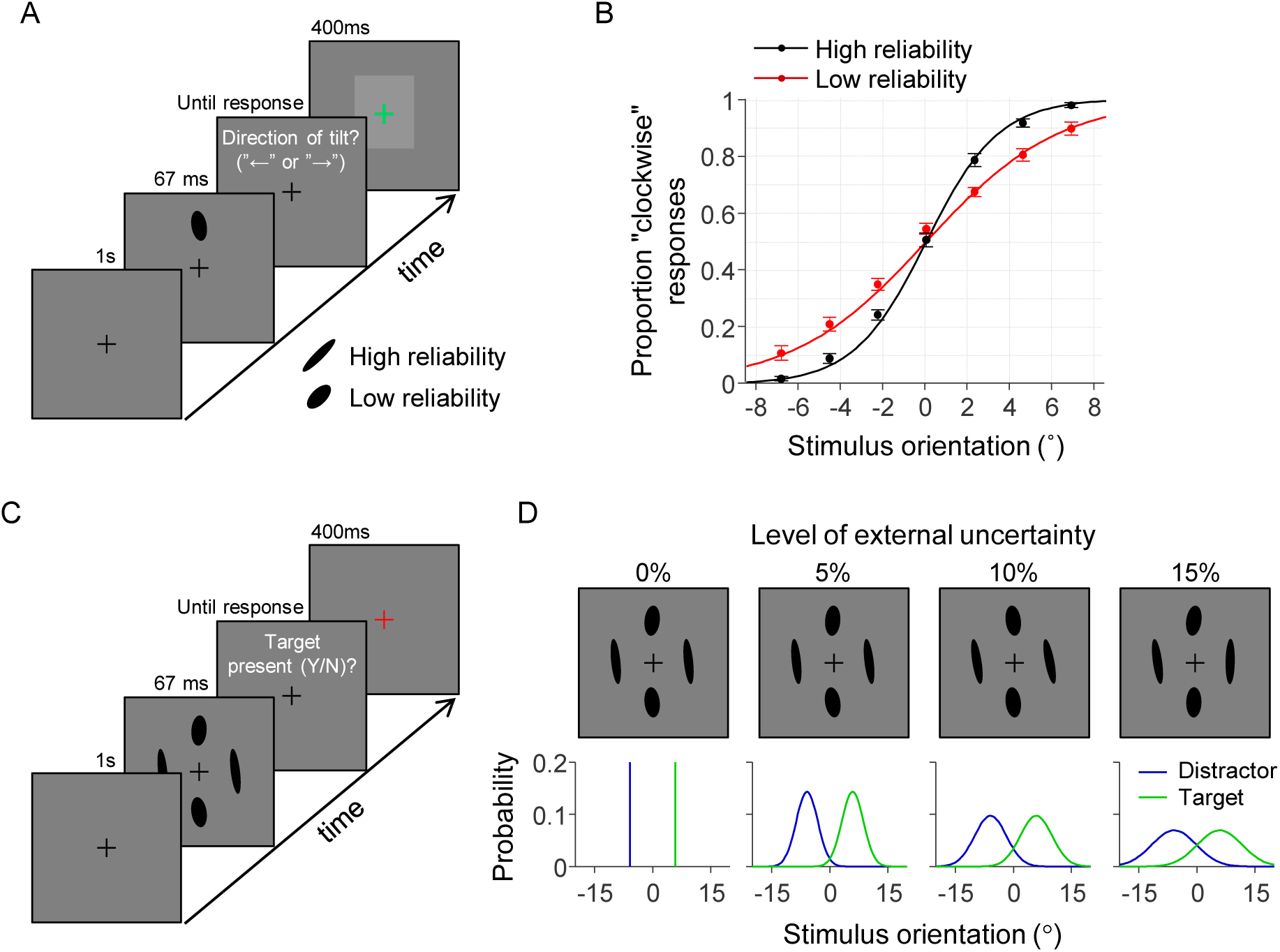
Experimental design. (A) Illustration of a trial in the discrimination task. Subjects reported on each trial the tilt direction of a single ellipse (“clockwise” or “counterclockwise” relative to vertical). The elongation of the stimulus could take two values. We refer to the most elongated type of ellipse as a “high reliability” stimulus and the less elongated type as a “low reliability” stimulus. Feedback was provided by briefly turning the fixation cross red (error) or green (correct) after the response was given. (B) The subject-averaged data (filled circles) and model fits (curves) reveal that sensitivity was higher for stimuli with high reliability (black) compared to those with low reliability (red). Error bars represent 1 s.e.m. (C) Illustration of a trial in the visual search task with brief stimulus presentation time. (D) Top: examples of target-present displays under the four different levels of external uncertainty. Bottom: distributions from which the stimuli in the example displays were drawn. In all four examples, the ellipse at the “north” location is a target and the other three are distractors.

### General procedure

Each subject completed multiple experimental sessions that lasted about one hour each. At the start of the first session, they received general information about the experiment. Thereafter, they performed a discrimination task (Fig. 1A) followed by one condition of the visual search task. In the remaining sessions, they only performed the visual search task (Fig 1C). We created eight conditions for the visual search task by using a 2×4 factorial design (Table 1). The factors specify the stimulus presentation time (short *vs.* unlimited) and the level of external uncertainty (none, 5%, 10%, and 15%; explained below). Different groups of subjects performed different subsets of these conditions.

**Table 1.** Overview of visual search task conditions and experimental subject groups. Each group consisted of 10 subjects. The condition with unlimited stimulus time and no external uncertainty was excluded from the experiment, because subjects are expected to perform 100% correct on it.

|  |  | Level of external uncertainty |  |  |  |
| --- | --- | --- | --- | --- | --- |
|  |  | None | 5% | 10% | 15% |
| Stimulus display time | Short (67 ms) | A<br>Group 1 | B<br>Group 2 | C<br>Group 1 | D<br>Group 3 |
|  | Unlimited | - | E<br>Group 2 | F<br>Group 1 | G<br>Group 3 |

### Discrimination task

On each trial, the subject was presented with a single ellipse (67 ms) and reported whether it was tilted clockwise or counterclockwise with respect to vertical (Fig. 1A). Trial-to-trial feedback was provided by briefly turning the fixation cross in the inter-trial screen green (correct) or red (incorrect). The elongation of the stimulus was 0.80 on half of the trials (“low reliability”) and 0.94 on the other half (“high reliability”), randomly intermixed. The stimulus location was randomly drawn on each trial from the set of four cardinal locations (“north”, “east”, “south”, and “west”). On the first 20 trials, the orientation of the stimulus was drawn from a uniform distribution on the range −5° to +5°. In the remaining trials, a cumulative Gaussian was fitted to the data collected thus far and the orientation for the next trial was then randomly drawn from the domain corresponding to the 55-95% correct range. This adaptive procedure increased the information obtained from each trial by reducing the number of extremely easy and difficult trials. Subjects completed 500 trials of this task.

### Visual search without external uncertainty (condition A)

In this condition, subjects were on each trial presented with four oriented ellipses. On half of the trials, all ellipses were distractors. On the other half, three ellipses were distractors and one was a target. The task was to report whether a target was present. Targets were tilted μ_target_ degrees in clockwise direction from vertical and distractors were tilted μ_target_ degrees in *counterclockwise* direction. The value of *µ*_target_ was customized for each subject (Table 2) such that an optimal observer with sensory-noise levels equal to the ones estimated from the subject’s discrimination-task data had a predicted accuracy of 85% correct (averaged over trials with different combinations of low and high reliability stimuli). Stimulus display time was 67 ms and each stimulus was presented with an ellipse elongation of either 0.80 (“low reliability”) or 0.94 (“high reliability”). On each trial, the number of high-reliability stimuli was drawn from a uniform distribution on integers 0 to 4 and reliability values were then randomly distributed across the four stimuli. The four stimuli always appeared at the four cardinal locations (“north”, “east”, “south”, and “west”). Feedback was provided in the same way as in the discrimination task. The task consisted of 1500 trials divided equally over 12 blocks with short forced breaks between blocks.

**Table 2.** Estimated sensory noise levels in the discrimination task 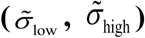 and the customized experimental parameters (μ_target_, *σ*_external_) in the visual search task.

| Level of external uncertainty (%) | Subj ID | $\tilde{\sigma}_{\text{low}}$ (°) | $\tilde{\sigma}_{\text{high}}$ (°) | $\mu_{\text{target}}$ (°) | $\sigma_{\text{external}}$ (°) |
| --- | --- | --- | --- | --- | --- |
| 0 | 1 | 7.1 | 4.6 | 8.0 | 0 |
| 0 | 2 | 5.5 | 2.0 | 5.0 | 0 |
| 0 | 3 | 6.4 | 2.3 | 5.8 | 0 |
| 0 | 4 | 3.8 | 1.8 | 3.8 | 0 |
| 0 | 5 | 4.6 | 2.2 | 4.5 | 0 |
| 0 | 6 | 4.0 | 3.3 | 5.0 | 0 |
| 0 | 7 | 3.1 | 1.3 | 2.9 | 0 |
| 0 | 8 | 6.8 | 2.8 | 6.4 | 0 |
| 0 | 9 | 3.3 | 2.4 | 3.9 | 0 |
| 0 | 10 | 4.5 | 3.0 | 5.1 | 0 |
| 5 | 11 | 4.4 | 3.4 | 5.3 | 2.4 |
| 5 | 12 | 3.7 | 3.0 | 4.5 | 2.1 |
| 5 | 13 | 4.2 | 2.6 | 4.6 | 2.1 |
| 5 | 14 | 3.9 | 2.2 | 4.1 | 1.9 |
| 5 | 15 | 4.3 | 3.1 | 4.9 | 2.3 |
| 5 | 16 | 6.5 | 2.4 | 5.9 | 2.8 |
| 5 | 17 | 6.2 | 3.3 | 6.4 | 2.9 |
| 5 | 18 | 3.8 | 2.0 | 3.9 | 1.8 |
| 5 | 19 | 5.1 | 3.0 | 5.4 | 2.5 |
| 5 | 20 | 4.6 | 2.6 | 4.8 | 2.2 |
| 10 | 1 | 7.1 | 4.6 | 8.0 | 5.6 |
| 10 | 2 | 5.5 | 2.0 | 5.0 | 3.4 |
| 10 | 3 | 6.4 | 2.3 | 5.8 | 4.1 |
| 10 | 4 | 3.8 | 1.8 | 3.8 | 2.6 |
| 10 | 5 | 4.6 | 2.2 | 4.5 | 3.1 |
| 10 | 6 | 4.0 | 3.3 | 5.0 | 3.4 |
| 10 | 7 | 3.1 | 1.3 | 2.9 | 2.1 |
| 10 | 8 | 6.8 | 2.8 | 6.4 | 4.3 |
| 10 | 9 | 3.3 | 2.4 | 3.9 | 2.7 |
| 10 | 10 | 4.5 | 3.0 | 5.1 | 3.4 |
| 15 | 21 | 6.2 | 2.6 | 5.9 | 5.8 |
| 15 | 22 | 7.7 | 2.4 | 6.8 | 6.7 |
| 15 | 23 | 6.9 | 4.2 | 7.5 | 7.1 |
| 15 | 24 | 6.7 | 4.8 | 7.9 | 7.5 |
| 15 | 25 | 7.5 | 4.5 | 8.2 | 7.8 |
| 15 | 26 | 5.1 | 3.6 | 5.8 | 5.7 |
| 15 | 27 | 4.1 | 2.3 | 4.3 | 4.3 |
| 15 | 28 | 8.1 | 3.3 | 7.7 | 7.4 |
| 15 | 29 | 7.2 | 5.2 | 8.5 | 8.1 |
| 15 | 30 | 7.1 | 3.3 | 6.9 | 6.8 |

### Visual search with external uncertainty and short display time (conditions B-D)

The three visual search conditions with external uncertainty and short display time were identical to condition A, except that the orientations of the target and distractors were not fixed, but instead drawn from partly overlapping Gaussian distributions (Fig. 1D). These distributions had means *µ*_target_ and −*µ*_target_ (see above), respectively, and a standard deviation *σ*_external_. The value of *σ*_external_ was customized for each subject (Table 2) such that the accuracy of an optimal observer would drop by 5, 10, or 15% compared to the same condition with *σ*_external_=0 (no external uncertainty). We refer to these percentages as *levels of external uncertainty.* Subjects completed 1500 trials divided equally over 12 blocks with short forced breaks between blocks.

### Visual search with external uncertainty and unlimited display time (conditions E-G)

These three conditions were identical to conditions B-D, except for the following two differences. First, stimuli were presented with an ellipse elongation of 0.97 and stayed on the screen until a response was provided, such that the sensory noise levels were reduced to a presumably negligible level. Second, this condition contained 500 instead of 1500 trials. Each subject completed this condition before the equivalent condition with short display times.

### Statistical analyses

All statistical tests were performed using the JASP software package [35]. Besides *p* values we also report Bayes factors, which specify the ratio between how likely the data are under one hypothesis (e.g., the null hypothesis) compared to how likely they are under an alternative hypothesis. An advantage of Bayes factors is that they can be used to both reject and support a hypothesis, whereas *p* values can only reject. All reported Bayes factors were computed using the default settings for the effect size priors (Cauchy scale parameter = 0.707; *r* scale for fixed effects = 0.5).

## MODELS

This section describes the models that we will fit to the data from the visual search task. An overview of these models is presented in Table 3.

**Table 3.** Overview of models and their free parameters for the visual search task with short display time. Parameters *σ*_low_ and *σ*_high_ only exist when the models are applied to conditions with short display time; in conditions with unlimited display time, the sensory noise level is fixed to a prespecified value (explained in Results).

| Model ID | Label | Free parameters | # |
| --- | --- | --- | --- |
| 1 | Flawless Bayesian | $\sigma_{\text{low}}, \sigma_{\text{high}}, \lambda$ | 3 |
| 2 | Imperfect Bayesian | $\sigma_{\text{low}}, \sigma_{\text{high}}, \lambda, \mu_{\text{late}}, \sigma_{\text{late}}$ | 5 |
| 3 | Ignorant Bayesian | $\sigma_{\text{low}}, \sigma_{\text{high}}, \lambda, \sigma_{\text{single}}$ | 4 |
| 4 | Imperfect ignorant Bayesian | $\sigma_{\text{low}}, \sigma_{\text{high}}, \lambda, \sigma_{\text{single}}, \mu_{\text{late}}, \sigma_{\text{late}}$ | 6 |
| 5 | Maximum of observations | $\sigma_{\text{low}}, \sigma_{\text{high}}, \lambda, c$ | 4 |
| 6 | Minimum deviation from target | $\sigma_{\text{low}}, \sigma_{\text{high}}, \lambda, c$ | 4 |
| 7 | Minkowski distance from target | $\sigma_{\text{low}}, \sigma_{\text{high}}, \lambda, c, \beta$ | 5 |
| 8 | Mean of observations | $\sigma_{\text{low}}, \sigma_{\text{high}}, \lambda, c$ | 4 |
| 9 | Variance of observations | $\sigma_{\text{low}}, \sigma_{\text{high}}, \lambda, c$ | 4 |
| 10 | Imperfect maximum-of-observations | $\sigma_{\text{low}}, \sigma_{\text{high}}, \lambda, c, \sigma_{\text{late}}$ | 5 |
| 11 | Imperfect minimum-deviation-from-target | $\sigma_{\text{low}}, \sigma_{\text{high}}, \lambda, c, \sigma_{\text{late}}$ | 5 |
| 12 | Imperfect Minkowski-distance-from-target | $\sigma_{\text{low}}, \sigma_{\text{high}}, \lambda, c, \beta, \sigma_{\text{late}}$ | 6 |
| 13 | Imperfect mean-of-observations | $\sigma_{\text{low}}, \sigma_{\text{high}}, \lambda, c, \sigma_{\text{late}}$ | 5 |
| 14 | Imperfect variance-of-observations | $\sigma_{\text{low}}, \sigma_{\text{high}}, \lambda, c, \sigma_{\text{late}}$ | 5 |

### Optimal Bayesian decision variable

Before introducing the models, we derive the Bayesian decision variable for our visual search task. We denote target presence by a binary variable *T* (0=absent, 1=present), set size by *N*, the stimulus values by **s**={*s*_1_, *s*_2_, …, *s_N_*}, and the observer’s noisy observations of the stimulus values by **x**={*x*_1_, *x*_2_, …, *x*_*N*_}. We make the common assumption that each stimulus observation, *x*_*i*_, is corrupted by zero-mean Gaussian noise, i.e., *x*_*i*_=*s*_*i*_+ε, where ε is a Gaussian random variable with a mean of zero. The standard deviation of this noise distribution, denoted *σ*_*i*_, is assumed to depend on the reliability of the stimulus, which in our experiment differed across locations (low *vs.* high reliability). The Bayesian observer reports “target present” if the posterior probability of target presence exceeds that of target absence, *p*(*T*=1|**x**)>*p*(*T*=0|**x**). This strategy is equivalent to reporting “target present” if the log posterior ratio exceeds 0,

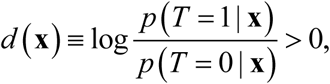

where *d*(**x**) is referred to as the global decision variable. Under the generative model for our task (S1 Figure) this evaluates to

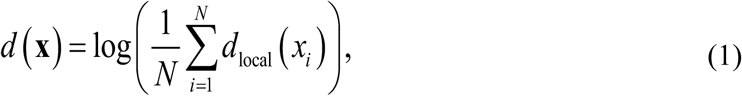

where

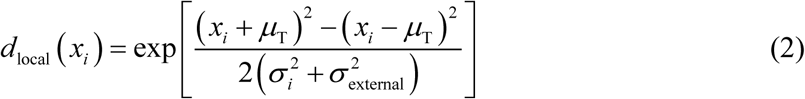

is referred to as the local decision variable (see S1 Appendix for a derivation). Hence, the optimal decision variable is the log of an average of local decision variables, each of which represents the evidence (posterior ratio) for target presence: *d*_local_(*x*_*i*_)<1 is evidence for a distractor at location *i* and *d*_local_(*x*_*i*_)>1 is evidence for a target; a value of exactly 1 represents equal evidence for both options. We mentioned earlier that optimal observers weight each cue by its reliability. In Eq. (2), this weighting occurs through sensory noise levels *σ*_*i*_: the larger *σ*_*i*_, the closer the local evidence associated to stimulus *x*_*i*_ is to 1.

#### Note about the sensory noise distribution

Since our stimulus domain is circular, the choice of a non-circular (Gaussian) noise distribution may seem poorly motivated. A theoretically better choice would have been to use a Von Mises distribution, as we have done in previous work (e.g., [15,19]). However, the sensory noise levels in our study are relatively low, in which case the Gaussian is a near-perfect approximation to the Von Mises. Because of its analytical and computational convenience, we decided to use a Gaussian rather than Von Mises noise distribution.

### Model 1: The flawless Bayesian

The first model that we consider is the Bayesian observer without any imperfections beyond sensory noise. This observer – which we refer to as the “flawless Bayesian” – is assumed to have perfect knowledge of the statistical structure of the task and to use Eq. (1) to compute its decision variable. Moreover, the flawless Bayesian is assumed to compute without error. The model’s only free parameters are the sensory noise levels *σ*_*i*_. In conditions with unlimited display time, we fix *σ*_*i*_ either to 0 (no noise) or to a value obtained from a control experiment (explained in Results). In conditions with short display time, we fit *σ*_*i*_ separately for stimuli with low reliability (*σ*_low_) and stimuli with high reliability (*σ*_high_). Heeding the concern that an excess of flexibility in optimal models can make suboptimal behavior look optimal [23], we constrain these parameters by imposing prior distributions on their values (see S1 Appendix). Moreover, we refrain from adding a bias parameter to this model, for two reasons. First, while many previous studies – including some of our own (e.g. [19,21]) – have not considered it problematic to allow for bias when testing for optimality, being biased is strictly speaking a violation of optimality. Second, and more importantly, a response bias can be confounded with biases caused by other, less obvious kinds of suboptimalities, as we will explain in our presentation of Model 2.

### Model 2: The imperfect Bayesian

Our second model is a Bayesian observer with imperfections in the computation of the decision variable. Such imperfections may produce suboptimalities in performance and could be caused by many different factors, such as noise in the neural mechanisms that compute the decision variable, incomplete knowledge of the statistical structure of the task, uncertainty about the experimental parameters, and suboptimal cue weighting. To get an idea of how computational imperfections affect a Bayesian observer’s decisions, we perform simulations with imperfect variants of Model 1. The imperfections in these variants create errors in the model’s decision variable, as compared to the decision variable of the flawless Bayesian observer. We simulate a large number of trials and find that for all tested imperfections, the distribution of this error is reasonably well approximated by a Gaussian distribution (Fig. 2). Importantly, the mean of this Gaussian is not always zero, which indicates that computational imperfections may produce a systematic error in the decision variable, i.e., a bias. Since this computational bias is indistinguishable from a simple response bias, the two can easily be confounded, which is the main reason why we did not include a response bias in the flawless Bayesian model.

**Fig. 2.**
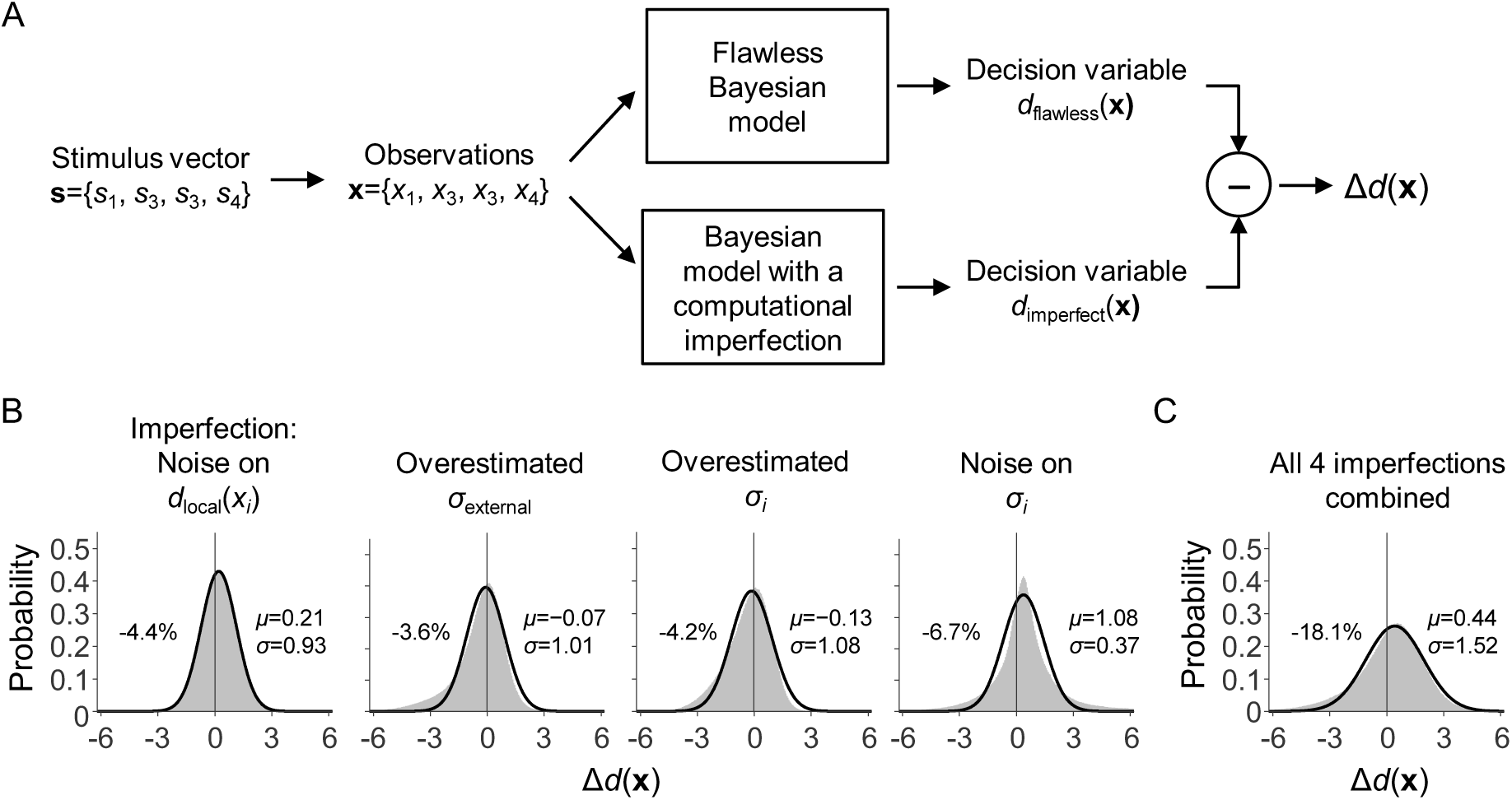
Simulated effects of four computational imperfections. (A) Schematic illustration of a single trial in the simulation that was aimed at assessing how computational imperfections affect the optimal observer’s decision variable. On each trial, a stimulus set s and stimulus observations x were drawn from the generative model for the visual search task with 10% external uncertainty. Next, x was provided as input to the Flawless Bayesian model and to a variant of this model with a computational imperfection (e.g., a wrong belief about experimental parameter *σ*_external_). Both models produce a decision variable, *d*(x). We denote the difference between these two decision variables by Δ*d*(x), which can be thought of as a computational error. A total of 1 million trials was simulated using four different types of computational imperfection: (1) Gaussian noise on the local decision variables; (2) an overestimated value of *σ*_external_; (3) overestimated values of *σ*_low_ and *σ*_high_; (4) item-to-item and trial-to-trial noise on *σ*_low_ and *σ*_high_. (B) The distribution of Δ*d*(x) under each simulated computational imperfection (gray areas). In all four cases, this distribution is reasonably well approximated by a Gaussian distribution (black curves). The percentages indicate the accuracy loss caused by the computational imperfection; parameters μ and *σ* indicate the mean and standard deviation of the Gaussian fitted to each distribution. (C) The distribution of Δ*d*(x) in a model that contains all four tested imperfections simultaneously.

The finding that different kinds of suboptimality produce similar errors in the decision variable implies that it will be difficult to distinguish between them in model comparison. However, the upside of this similarity is that it allows us to test for computational imprecisions in a rather *general* way: instead of implementing a separate model for each possible computational imperfection, we can test for a range of different imperfections by using a single model with Gaussian noise on the optimal decision variable. We implement this “imperfect Bayesian” model by adding a noise term η to Eq. (1),

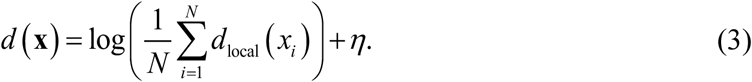

We denote the mean (bias) and standard deviation of this “late” noise by μ_late_ and *σ*_late_, respectively, which are fitted as free parameters.

### Models 3 and 4: The ignorant Bayesian

The first two models weight each stimulus by its reliability, which is a hallmark of Bayesian observers. Model 3 is a variant that ignores differences in cue reliabilities and instead weighs them equally. In this model, Eq. (2) is replaced with

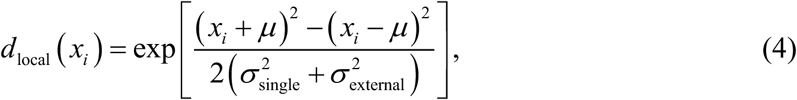

where *σ*_single_ is a free parameter that determines the weight assigned to every stimulus. For lack of a better term, we refer to this model as the “ignorant Bayesian”. Model 4 is a variant of this model in which we add computational imperfections in the same way as in Model 2, i.e., by adding biased Gaussian noise to the global decision variable.

### Models 5-9: Heuristic models

In Models 1 – 4, decisions were made based on the optimal decision variable or an impoverished variant of it. We next introduce five models with heuristic decision strategies. The first of them uses the maximum-of-output or “max” rule, which has its origin in signal detection theory [36] and is a commonly used heuristic in models of visual search (e.g., [16,37–39]). In the present task, the rationale is that since target orientations are on average larger than distractor orientations, one might perform well by reporting “target present” whenever the maximum observation, *x*_*i*_, exceeds some threshold, *c*. The decision variable of the Max model is thus simply the maximum stimulus observation,

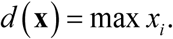

The next two heuristic models make decisions based on how much the stimulus observations deviate from the expected target value. Model 6 uses the minimum deviation as its decision variable,

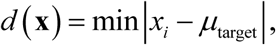

which again is compared with a decision criterion *c*. Similarly, Model 7 uses the Minkowski distance between any stimulus observation and the expected target value as its decision variable,

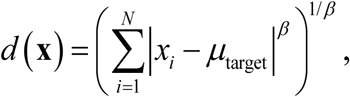

where *β* is a free parameter. The next and final two heuristics are inspired by previous findings that the visual system represents summary statistics of the stimuli that it observes, including their mean and variance [40–42]. These statistics could be used to solve detection tasks of the kind used in our experiment, where both the mean and variance of the stimulus observations are expected to be larger on trials with a target compared to trials without a target. Therefore, Model 8 uses the mean of observations as the decision variable,

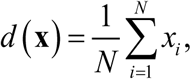

and Model 9 uses the variance,

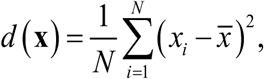

where 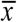 is the average of the stimulus observations.

#### Free parameters

Just as in the Bayesian models, sensory noise levels *σ*_low_ and *σ*_high_ are fitted as free parameters. In addition, since heuristic strategies do not dictate the value of the decision criterion, *c*, it is fitted as a free parameter as well (in Bayesian models, the decision criterion is 0 by design). The Minkowski model has an additional free parameter *β*. We fit all these parameters in an entirely unconstrained way. This means that we give more flexibility to the heuristic models than to the Bayesian models, in which parameters are constrained by imposing prior distributions. This way, we ensure that if we find evidence for Bayesian models, then it is unlikely to be due to them being more flexible than alternative models.

### Models 10-14: Imperfect heuristic models

The final five models that we consider are imperfect variants of the heuristic models. In these models, the decision variable is corrupted in the same way as in the imperfect Bayesian models. However, since bias in heuristic models is already captured in the criterion value, *c*, we fix μ_late_ to 0 and only fit *σ*_late_ as a free parameter.

### Lapse rate

Models of perceptual decision-making tasks often include a lapse rate to account for random guesses caused by attentional lapses. In such models, it is assumed that responses on some of the trials were the result of guessing rather than a decision strategy. The lapse rate parameter specifies the estimated proportion of guessing trials. If we do not include a lapse rate in our models, then we run the risk of underestimating how good the subjects’ decision strategies were, because guessing behavior can then only be accounted for as suboptimalities in their decision strategies. On the other hand, if we *do* include a lapse rate, then we give models a possibility to explain away decision suboptimalities as lapses, which brings along the opposite risk: we might *overestimate* how good subjects’ decision strategies were. In an attempt to minimize both risks, we include a lapse rate in all models, but in the Bayesian models we constrain this parameter by imposing a prior distribution on its values (see S1 Appendix).

### Model fitting and model comparison

We use an adaptive Bayesian optimization method [43] to find maximum-likelihood estimates of model parameters, at the level of individual subjects. Model evidence is measured as the Akaike Information Criterion [44] and interpreted using the rules of thumb provided by Burnham & Anderson [45]. We performed a model recovery analysis [46] to verify that the models make sufficiently diverging predictions to distinguish them in a model comparison (see S2 Figure).

## RESULTS

### Discrimination task

Under the assumption that stimulus observations are corrupted by Gaussian noise, the predicted proportion of “clockwise” responses in the discrimination task is a cumulative Gaussian function of stimulus orientation. We refer to the standard deviation of this Gaussian as the sensory noise level. To verify that differences in stimulus elongation caused differences in sensory noise levels, we fitted two cumulative Gaussian models to the data. In the first model, the noise level is independent of ellipse elongation and fitted as a single free parameter. In the second model, the sensory noise levels are fitted as separate parameters for the low-and high-reliability stimuli, which we denote by 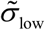 and 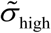, respectively. The second model accounts well for the data (Fig. 1B) and model comparison favors this model for every subject (ΔAIC range: 0.50 to 22.3; mean±sem: 8.6±1.3). Moreover, for every subject the estimated noise level is higher for the low-reliability stimulus than for the high-reliability stimulus (Table 2). Hence, the stimulus-reliability manipulation works as intended. We use noise estimates 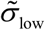 and 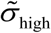 to customize the target and distractor distributions in the visual search experiment (Table 2) and to constrain the Bayesian models fitted to the data from that experiment (S1 Appendix).

While previous studies (e.g., [47]) have reported that performance on discrimination tasks is sometimes better for stimuli at the vertical meridian (“north”/“south” locations) than for stimuli at the horizontal meridian (“east”/“west” locations), we do not find evidence for such an effect in the present experiment. Performance differed little across locations, ranging from 74.3±1.1% correct at the “east” location to 75.0±1.0% at the “north” location. A Bayesian one-way ANOVA provides strong evidence for the null hypothesis of there being no effect (BF_01_=20.5, *p*=.97).

### Visual search with unlimited display time

We assume for the moment that sensory noise in the visual search conditions with unlimited display time was negligible, i.e., *σ*_*i*_=0. Under this assumption, the stimulus observations are identical to the true stimulus values, **x**=**s**, which allows us to write the optimal decision variable, Eq. (1), directly as a function of **s**,

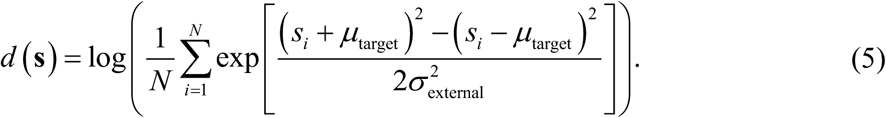

Since there are no unknowns in this equation, we can compute the optimal decision variable for each trial that was presented to a subject. The flawless Bayesian responds “target present” on each trial with *d*(**s**)>0 and “target absent” otherwise. Hence, if subjects are optimal, then their proportion of “target present” responses should be a step function of *d*(**s**), transitioning from 0 to 1 at *d*(**s**)=0. In all three conditions, subjects clearly deviate from this prediction (Fig. 3B, circles).

**Fig. 3.**
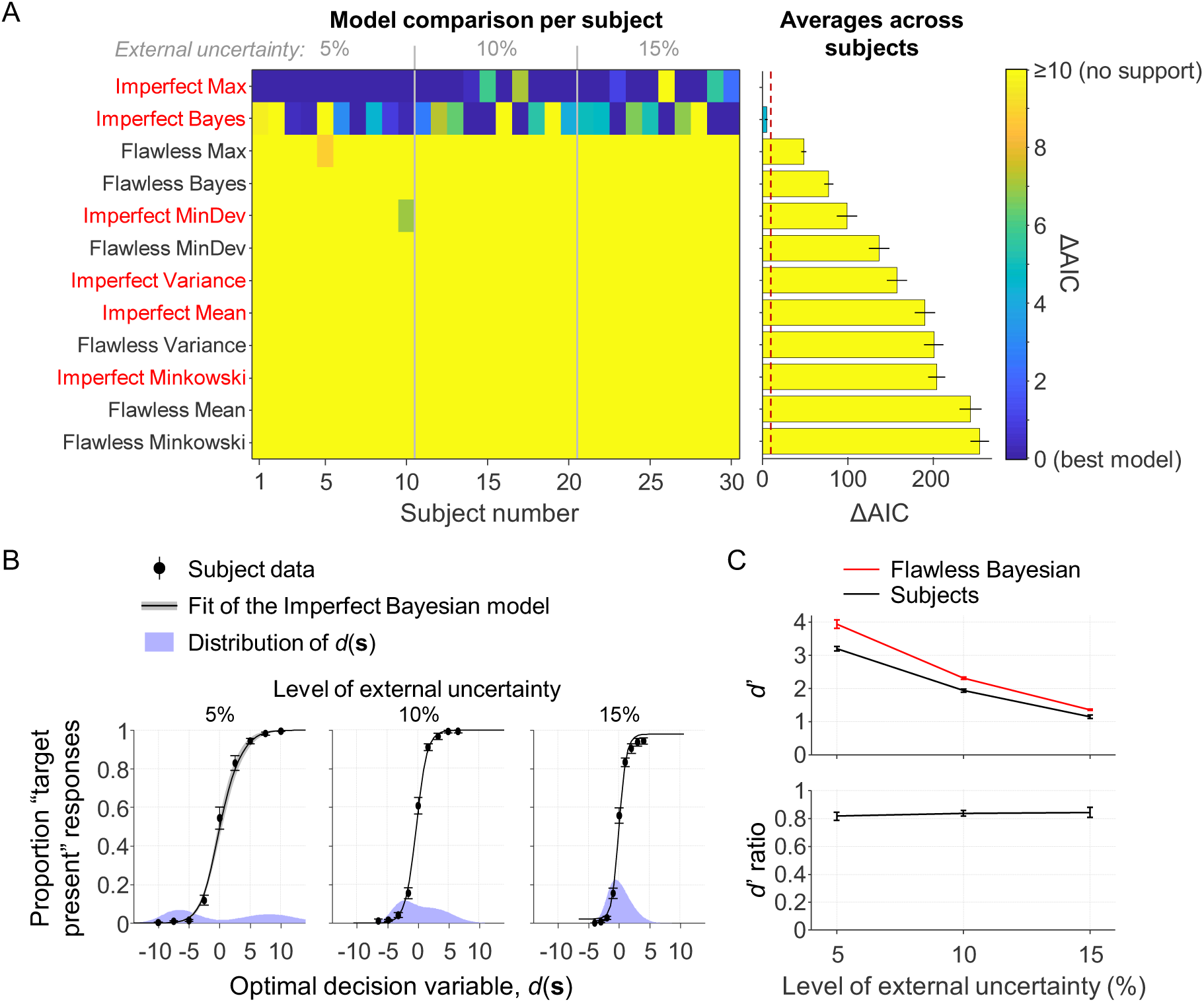
Results from the visual search conditions with unlimited display time. (A) Left: AIC-based model comparison at the level of single subjects. Each column is a subject and each row is a model. The best model for each subject is indicated in dark blue (ΔAIC=0). Right: Subject-averaged AIC values relative to the overall best model. The red dashed line indicates the ΔAIC≥10, which is interpreted as “no support”. (B) The subject data (black markers) are well accounted for by the “Imperfect Bayesian” and “Imperfect Max” models (black curves; the fits of both models are visually indistinguishable). Note that the distribution of *d*(**s**) (purple areas) becomes more concentrated around zero as the level of external uncertainty increases, due to the evidence generally being weaker in the tasks with more external uncertainty. (C) In all three conditions, the empirical *d*’ values (black) are lower than the values predicted by the Flawless Bayesian model (red). The average ratio between the *d*’ values is 0.834±0.017.

#### Model fits

To obtain insight into the possible nature of this apparent suboptimality, we fit the models listed in Table 3 to the individual datasets. Assuming that sensory noise is absent in these conditions, we set *σ*_i_=0 for all stimuli. Models 3 and 4 are excluded from the analysis, because they are identical to Models 1 and 2, respectively, when there is no sensory noise. Model comparison (Fig. 3A) selects the Imperfect Max model as the preferred model, closely followed by the Imperfect Bayesian model (ΔAIC=4.4±1.5). Both models account well for the data (Fig. 3B, curves) and all other models are rejected (ΔAIC≥48.0±3.1 relative to the selected model). The slight advantage of the Max model in model comparison seems to be entirely due to its flexibility in fitting the lapse rate parameter: when constraining this parameter in the same way as in the Bayesian model, the difference changes to ΔAIC=2.7±2.1 in favor of the Bayesian model.

We draw three conclusions from these model comparison results. First, the Max and Bayesian models are indistinguishable in these conditions (which was expected, as explained below). Second, the results provide strong evidence against the other four heuristics as well as against the flawless Bayesian. Third, whichever decision strategy was used, it seems that there were computational imprecisions in its execution. Model comparison using cross validation instead of AIC gives the same results and conclusion (S3 Figure).

#### Optimality index

While the above analysis suggests a deviation from optimality, it does not quantify the magnitude of this deviation. To estimate this magnitude, we introduce an optimality index *I* based on sensitivity indices,

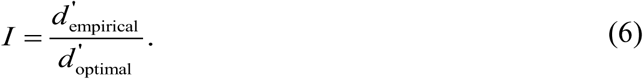

The numerator is the empirical sensitivity index, which we compute as the difference between the z-scores of the hit and false alarm rates in the subject data. The denominator is the sensitivity index of the optimal observer, which we compute in the same way, but based on simulated response data from the Flawless Bayesian model. In these simulations, we set the values of μ_target_ and *σ*_external_ to the subject’s customized values (Table 2) and lapse rate λ to the maximum-likelihood estimate of the best-fitting model.

The subject-averaged optimality index across all three conditions is 0.834±0.017 (Fig. 3C, bottom), which corresponds to a deviation of 16.6±1.7% from optimal. A Bayesian one-way ANOVA provides moderate evidence against the hypothesis that the optimality index depends on the level of external uncertainty (BF_01_=4.03; *p*=.79).

#### Accounting for possible sensory noise

Despite the unlimited display time, it is possible – and perhaps even likely – that there was still some noise in the subjects’ encoding of stimulus orientations. If that is the case, then our assumption *σ*_*i*_=0 was wrong and the above analysis will have underestimated the optimality index. Unfortunately, we cannot fit *σ*_*i*_ as a free parameter, because that creates identifiability problems in models with a *σ*_late_ parameter. Therefore, we instead estimate it using a separate experiment. This control experiment is identical to the discrimination experiment (Fig. 1A), except that the stimulus has an ellipse elongation of 0.97 and stays on the screen until a response is given. By fitting a cumulative Gaussian to the data from twelve (new) observers, we find an estimate *σ*_*i*_=0.875±0.097.

We fit the models again, but now with *σ*_*i*_ fixed to 0.875 instead of 0. Model comparison gives very similar results as before: the imperfect variants of the Max and Bayesian models are very close to each other (ΔAIC=5.0±1.5 in favor of the Max model) and none of the other models is supported (ΔAIC>47.8±8.2 relative to the best-fitting model). However, we now find a slightly higher optimality index, *I*=0.879±0.019, which corresponds to a 12.1±1.9% deviation from optimal. A Bayesian one-way ANOVA again suggests that there is no effect of the level of external uncertainty on the optimality index (BF_01_=2.40; *p*=.36).

### Visual search with short display times

Next, we fit the models to the data from the conditions with short display times. Model comparison (Fig. 4A) selects the Imperfect Bayesian as the preferred model and rejects all other models with large margins (ΔAIC≥19.6±4.0). This result is consistent with the results above, except that both Max models are now convincingly rejected. The main conclusion that we draw from this model comparison result is that subjects neither seem to behave optimally, nor do they seem to use a heuristic decision strategy. Instead, their decisions seem to be based on Bayesian principles that are implemented or executed imperfectly. Model comparison using cross-validation gives near-identical results (S3 Figure).

**Fig. 4.**
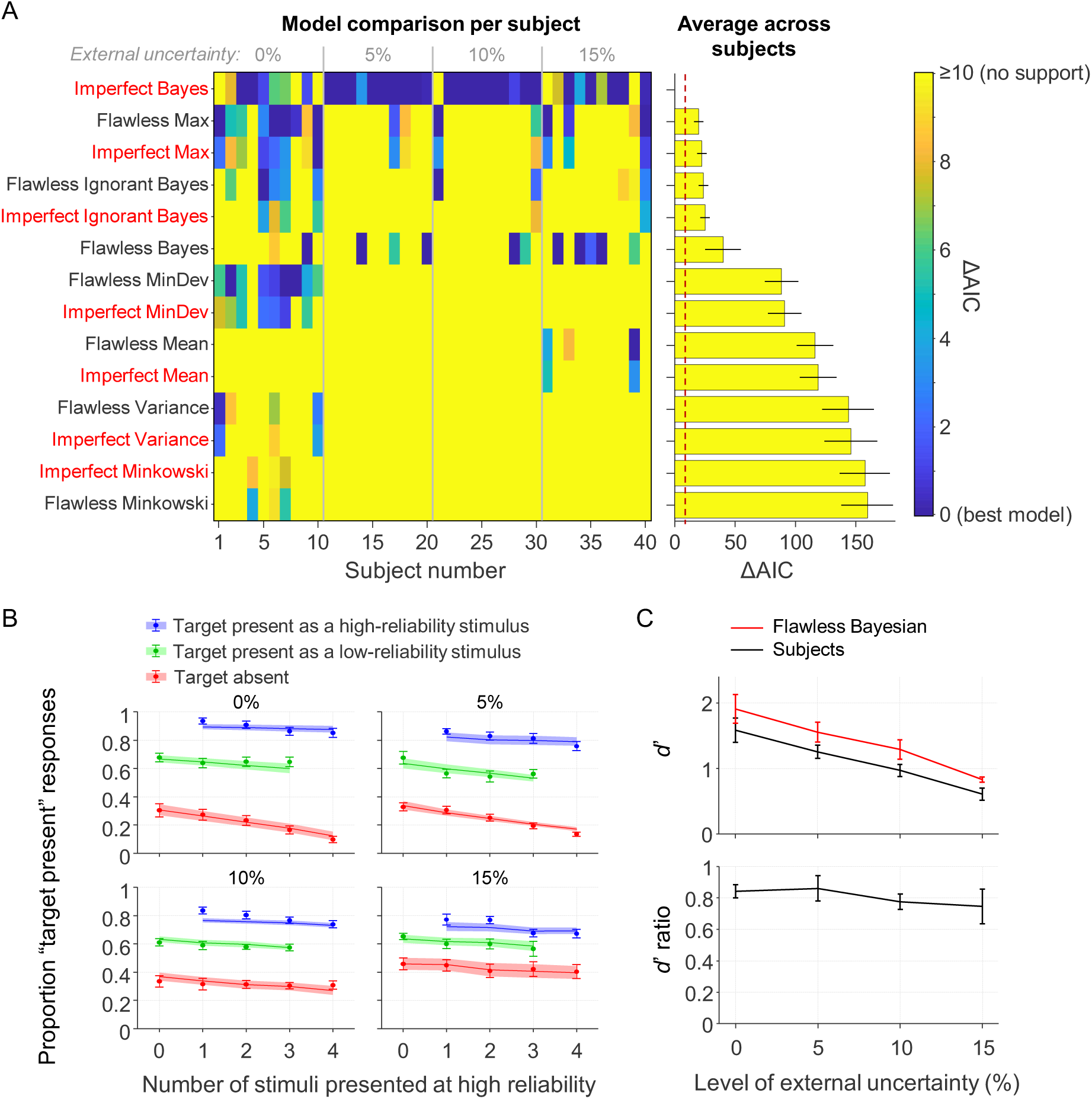
Results from the visual search conditions with short display time. (A) Left: AIC-based model comparison at the level of single subjects. Each column is a subject and each row is a model. The best model for each subject is indicated in dark blue (ΔAIC=0). Right: Subject-averaged AIC values relative to the overall best model. The red dashed line indicates the ΔAIC≥10, which is interpreted as “no support”. (B) The subject data (black markers) are well accounted for by the Imperfect Bayesian and Imperfect Max models (black curves; the fits of both models are visually indistinguishable). (C) In all three conditions, the empirical *d*’ values are lower than the values predicted by the Flawless Bayesian model.

The reason why the Max and Bayesian model were tied in the conditions with unlimited display time is that they make near-identical predictions when all stimuli have the same reliability, as we have observed in earlier work [48]. Therefore, it is important to use mixed-reliability designs when testing these two models against each other: while reliability-based weighting is an inherent property of Bayesian decision making, there is no natural way to incorporate such weighting in a Max model. Our finding that only a Bayesian model accounts well for data from mixed-reliability conditions strongly suggests that humans take stimulus reliability into account during perceptual decision making.

It is worth noting that it is unlikely that the superiority of the Imperfect Bayesian was due to it being overly flexible. First, it does not have more parameters than most of the heuristic models (Table 3). Second, while parameters in the heuristic models were entirely unconstrained, we imposed prior distribution on parameters of the Bayesian models. Third, a model recovery analysis (S2 Figure) showed that the Imperfect Bayesian is never selected when data are generated from one of the other 13 models.

#### Optimality index

We use the earlier introduced optimality index, Eq. (6), to estimate how much subjects deviate from optimality. We again use the Flawless Bayesian to compute 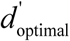, with *σ*_low_, *σ*_high_, and lapse rate λ set to the subject’s maximum-likelihood estimates in the best-fitting model. Averaged across all subjects in the conditions with brief display times, we find *I*=0.808±0.037, which corresponds to a 19.2±3.7% deviation from optimal performance (Fig. 4C). A one-way ANOVA reveals moderate evidence against an effect of the level of external uncertainty (BF_01_=5.05; *p*=.70). A two-way Bayesian ANOVA that also includes the optimality indices from the conditions with unlimited display time reveals moderate evidence against an effect of the level of internal uncertainty (BF_inclusion_=0.20; *p*=.47) and strong evidence against an effect of the level of external uncertainty (BF_inclusion_=0.07; *p*=.79). Averaged across all 7 experimental conditions, we find an optimality index of 0.819±0.022, which corresponds to an 18.1±2.2% deviation from optimality.

### Comparison with effects of sensory noise

The optimality indices reported above estimate how much performance was lost due to computational imprecisions. For comparison, we also estimate performance loss caused by sensory noise. To this end, we compute a variant of the earlier introduced optimality index, Eq. (6). In this variant, we “turn off” the sensory noise when computing 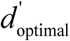, by fixing *σ*_*i*_ to 0. This new optimality index expresses empirical performance relative to an optimal observer without sensory noise. We refer to our original index as the “relative optimality” index and to this new index as the “absolute optimality” index [3]. The difference between these two indices gives an estimate of the amount of optimality loss due to sensory noise. To illustrate this, consider an example in which a subject has a relative optimality index *I*_relative_=0.80 and an absolute optimality index *I*_absolute_=0.70. In this example, the subject has an optimality loss of 0.20 when sensory noise is not considered to be a form of suboptimality and a loss of 0.30 when it is. We would in this case conclude that sensory noise accounted for 33.3% of the optimality loss (0.10 out of a total loss of 0.30) and computational imprecisions for 66.7% (0.20 out of 0.30).

When applying this method to the data from conditions with unlimited display time, we find that computational imperfections account for an estimated 92.6±3.8% of the performance loss and sensory noise for the remaining 7.4±3.8%. In conditions with short display time, we find that computational imperfections account for 27.0±5.1% of the performance loss and sensory noise for 73.0±5.1%. As expected, when sensory noise levels are low, performance loss is almost entirely attributed to computational imprecisions. Nevertheless, even in conditions with considerable levels of sensory noise, we estimate that almost a third of the performance loss was due to computational imprecisions.

### Analysis of parameter estimates

Next, we have a look at the best-fitting parameter estimates in the Imperfect Bayesian model. One-way ANOVAs suggest that there is an effect of the level of external uncertainty on both *σ*_low_ (BF_10_=9.06; *p*=.005) and *σ*_high_ (BF_10_=1.78; *p*=.042). Visual inspection of the parameter estimates (Fig. 5) reveals that this is mainly due to the condition with the highest level of external uncertainty, in which the sensory noise estimates are visibly higher than in the other conditions. However, the stimuli were extremely similar between the different conditions, which makes it implausible that there were large differences in sensory noise levels. Hence, despite our efforts to constrain these parameters, they may still have been overestimated in the condition with the highest level of external uncertainty. This means that we might have underestimated the magnitude of the deviation from optimality in that condition. For the two parameters that control the late noise distribution, we find neither an effect of the level of internal uncertainty (BF_inclusion_=0.25 for both μ_late_ and *σ*_late_) nor of the level of external uncertainty (μ_late_: BF_inclusion_=0.52; *σ*_late_: BF_inclusion_=0.10). Finally, for the lapse rate parameter we find evidence against an effect of the level of internal uncertainty (BF_inclusion_=0.56) and in favor of an effect of the level of external uncertainty (BF_inclusion_=1.22). However, the evidence for this effect is very weak and the estimated lapse rates are very small in all conditions, so we do not consider this finding to be of any significance.

**Fig. 5.**
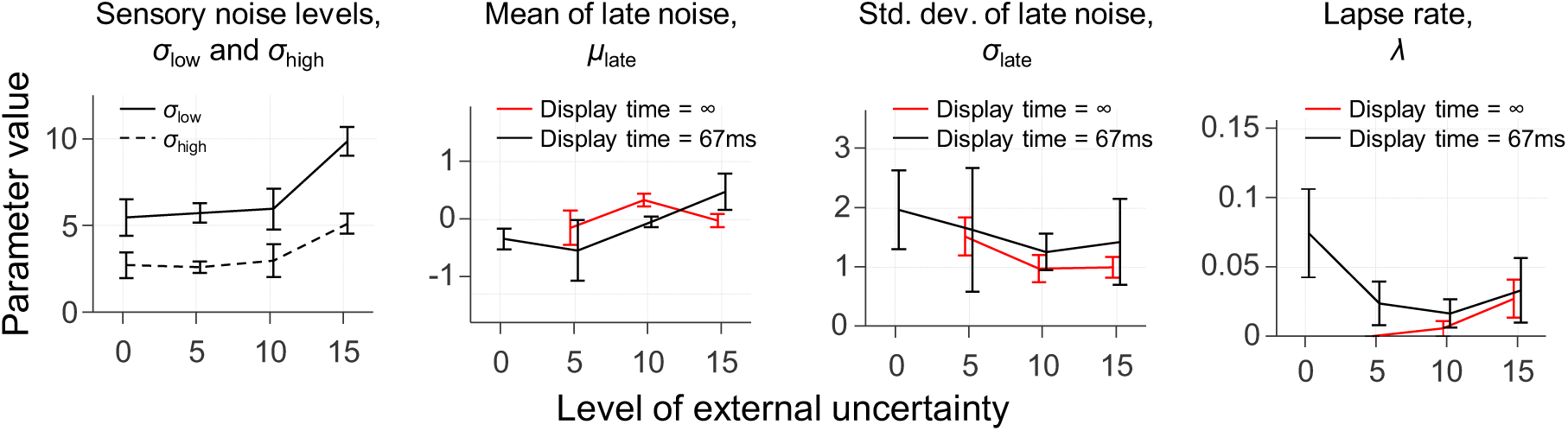
Maximum likelihood estimates of the parameters in the Imperfect Bayesian model. The reported parameters for conditions with unlimited display time were obtained with the model variant in which *σ*_*i*_ was fixed to 0.875.

### Reanalysis without pop-out trials

While the use of mixed reliabilities is a powerful way to test predictions that are unique to Bayesian models, it has the side effect that a stimulus may “pop out” when its reliability differs from that of all other stimuli. Stimuli that pop out may inadvertently draw attention and be given more weight, which would cause a suboptimality in performance, because the optimal weight is entirely determined by the reliability of a stimulus. We find that accuracy was slightly higher on trials in which the target popped out (72.5% correct) than on trials in which it did not (69.4% correct), which suggests that pop-out items indeed drew subjects’ attention. A t-test supports that there is a difference in accuracy between these two groups of trials (BF_10_=4.92; *p*=.008). To verify that the deviation from optimality in the conditions with short display time were not entirely caused by this pop-out effect, we fit the models again after filtering out pop-out trials. In this analysis, we thus only consider trials with 0, 2, or 4 high-reliability stimuli (60% of the data). Note that only a third of the trials in this modified dataset has mixed reliability.

As before, we find that model comparison selects the Imperfect Bayesian as the preferred model. However, the difference with the Max models is smaller now (Flawless Max: ΔAIC=7.0±2.2; Imperfect Max: ΔAIC=9.3±2.3; the difference with all other heuristic models is still large, ΔAIC≥48.8±7.9). This was to be expected, because we filtered out most of the mixed-reliability trials and we already established that the Max and Bayesian decision rules are indistinguishable on single-reliability data. When we constrain the parameters in the Max model in the same way as in the Bayesian models – which makes a fairer comparison – the Imperfect Bayesian outperforms both Max models with decent margins (Flawless Max: ΔAIC=10.3±2.5; Imperfect Max: ΔAIC=15.6±2.6). The optimality index in this analysis is 0.797±0.026, which is nearly identical to the value we obtained in the analysis that included all trials (0.808±0.037). Indeed, a t-test provides moderate evidence for the null hypothesis that there is no difference (BF_01_=3.58, *p*=.52). Altogether, our conclusions are largely the same under inclusion and exclusion of pop-out trials, which suggests that pop-out effects play a relatively minor role in explaining the identified suboptimalities.

### Reanalysis without a lapse rate

So far, we have included a lapse rate in all our models. To assess whether our conclusions would have been different if we had not included a lapse rate, we rerun all analyses with lapse rates fixed to 0. The model comparison results are very similar to the results reported above: in conditions with unlimited display time, the imperfect Max and Bayesian models are indistinguishable (ΔAIC=1.29±0.95 in favor of Bayes) and all other models are strongly rejected (ΔAIC≥96±12); in conditions with short display time, the imperfect Bayesian is selected as the preferred model and all other models are again strongly rejected (ΔAIC≥32.2±4.4). However, the optimality indices are now slightly lower: *I*=0.876±0.018 (13.3±1.8% deviation from optimality) in conditions with unlimited display time and *I*=0.796±0.037 (20.4±3.7% deviation from optimality) in conditions with brief display time. This was to be expected, because errors that were explained as lapses in our original analysis can now only be explained by suboptimalities in the decision strategy. As before, a two-way Bayesian ANOVA suggests that there is no effect of the level of external uncertainty (BF_inclusion_=0.167) nor of the level of internal uncertainty (BF_inclusion_=0.842) on the optimality index. Altogether, we conclude that removing the lapse rate from the models does not significantly change our conclusions.

### Reanalysis without constraints on model parameters

Finally, we check what happens to the results when we remove the constraints on the parameters of the Bayesian models, by refitting Models 1–4 without prior distributions on parameters *σ*_low_, *σ*_high_, and λ. The model comparison result is again very similar to our previous results: in conditions with unlimited display time, the imperfect Max and Bayesian models are indistinguishable (ΔAIC=1.4±1.1 in favor of Bayes) and all other models are strongly rejected (ΔAIC≥49.4±3.7); in conditions with short display time, the imperfect Bayesian model convincingly outperforms all other models, including the Max models (ΔAIC≥21.9±3.9). Also, we again find no evidence for an effect of the level of internal or external uncertainty on the optimality index (BF_inclusion_=0.56 and 0.19, respectively). However, the estimated deviation from optimality is now 11.7±2.1%, which is substantially lower than the 18.1±2.2% that we found with constrained parameter fits. This was to be expected, because without parameter constraints, models may explain away some of the computational suboptimalities by overestimating the lapse rate and/or sensory noise levels. Indeed, the average estimated lapse rate is now 13.7±2.2%, compared to 3.7±1.1% in the constrained fits (BF_+0_=582; *p*<.001). For some subjects the estimated lapse rate is now even over 50%, which seems unrealistically high. Hence, it appears that lapse rates are overestimated in the unconstrained fit. The estimated sensory noise levels, on the other hand, are very similar to the estimates obtained with the constrained fitting method (*σ*_low_=6.58±0.55 *vs.* 6.75±0.54; *σ*_high_=2.67±0.38 *vs.* 3.35±0.37). Indeed, a t-test supports the hypothesis that there is no difference (BF_01_=4.43; *p*=0.44). We speculate that the richness of data from mixed-reliability experiments is itself a sufficient constraint on the parameter values, in particular when subjects use a decision strategy that is sensitive to reliability differences between stimuli.

### Summary of results

A summary of the results from all analyses is presented in Table 4.

**Table 4.**
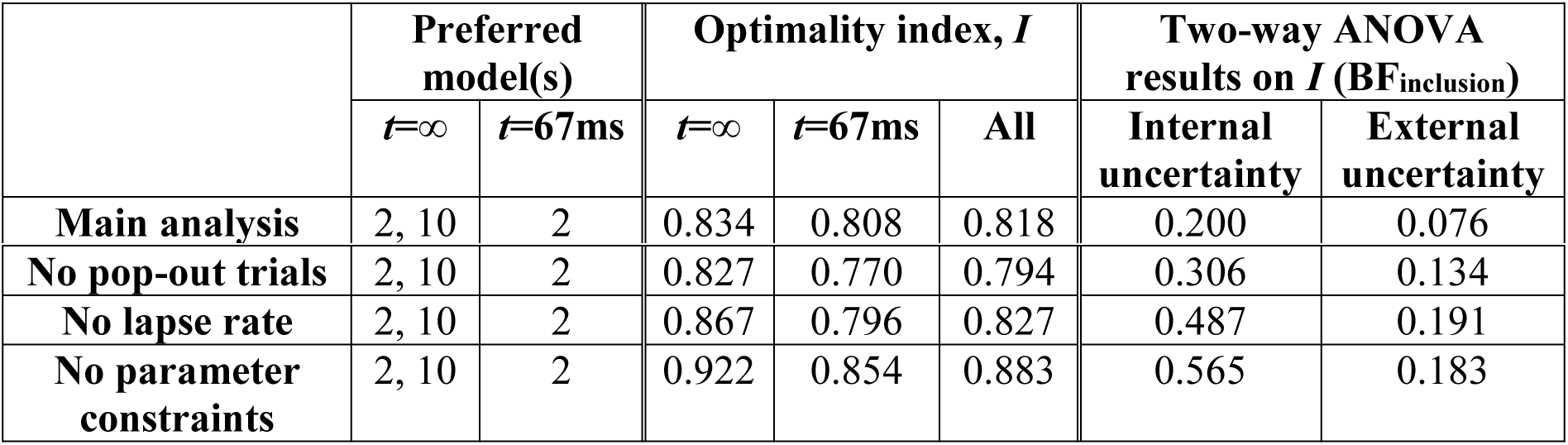
Summary of results across the four analyses. Models 2 and 10 are the Imperfect Bayesian and Imperfect Max models, respectively. Bayes factor BF_inclusion_ indicates whether there is evidence for an effect of internal or external uncertainty on the optimality index. All Bayes factors are smaller than 1, indicating evidence *against* an effect.

## DISCUSSION

In this study we re-examined optimality of human perception by using a standard visual search task. In contrast to previous claims that humans perform near-optimally on this task [13–16], we found no support for the Flawless Bayesian model. More specifically, we estimated that empirical performance deviated on average 18.1% from optimal performance. Interestingly, the estimated magnitude of this deviation did not depend on the level of internal uncertainty, nor on the level of external uncertainty. This stability may be a sign that the estimates were accurate, which would mean that our method successfully dissociated computational sources of suboptimality from sensory sources. Our data are best described by a model that is based on Bayesian principles, but with imperfections in the implementation of these principles. We believe that such “Imperfect Bayesian” models can provide a fruitful middle ground in the debate between Bayesian and anti-Bayesian views on human perception.

### Suboptimal behavior does not necessarily imply heuristic-based decision making

Deviations from optimality are often taken as evidence for heuristic decision making. However, this is not necessarily true: Bayesian observers can also be suboptimal. In particular, it has been argued that imprecisions in neural systems and the need to use deterministic approximations in complex computations may be the main reason why humans are unable to perform optimally on many tasks [27–31]. Such imperfections are orthogonal to the underlying decision strategy, because they may apply to both Bayesian and heuristic decision strategies. However, most previous work has only compared models with the optimal decision strategy against models with heuristic strategies, without testing for computational imprecisions. Claims of optimality made in those works are probably too strong, because evidence for a Bayesian decision strategy does not imply optimality. To avoid such overly strong claims, we advocate using a factorial modeling approach by crossing the decision strategy (Bayesian *vs.* heuristic-based strategies) with the absence or presence of computational imprecisions. Such an approach can decompose suboptimality into two different sources: using a fundamentally wrong decision strategy and having imperfections in the execution of this strategy. Only evidence for Bayesian decision-making without imprecisions should be considered as evidence for optimal behavior.

### Decomposing source of suboptimality

It has recently been argued that instead of focusing on the binary question whether or not a particular behavior is optimal, it would be more fruitful to start building process models that precisely characterize the sources that make humans prone to errors [23]. The approach that we took here can be seen as a step in this direction, as it aims at distinguishing between different kinds of suboptimality and quantifying the amount of performance loss caused by each of them. A similar approach was recently developed by Drugowitsch and colleagues [27], who examined sources of suboptimality in a visual categorization task. They estimated that about 90% of the performance loss was caused by imprecisions in mental inference and the remaining 10% by stochasticity in sensory input and response selection. In our visual search task, we found a numerically similar contribution of computational imprecisions in the conditions with unlimited display time (92.6%). However, in the conditions with brief display time, we found that only about a third of the optimality loss was due to computational imprecisions. This can be understood by considering that sensory noise levels were probably higher in our experiment, due to a difference in stimulus presentation time (67 ms to encode four stimuli in our study *vs.* 333 ms per stimulus in the study by Drugowitsch et al.). We tried to further decompose suboptimalities into more specific sources, such as “noise in the computation of local decision variables”, “incorrect knowledge of the experimental parameters”, and “suboptimal cue weighting”. However, as demonstrated by the simulation results presented in Fig. 2, different types of suboptimalities have near-identical effects on the response data, due to which we were unable to reliably distinguish between them using model comparison. Future studies may try to solve this model-identifiability problem by using experimental paradigms that provide a richer kind of behavioral data to further constrain the models (e.g., by collecting confidence ratings [49]).

### The importance of using mixed reliability designs

While reliability-based cue weighting is an inherent property of Bayesian observers, heuristic models do not have a natural way of taking reliability into account. Therefore, within-display manipulation of stimulus reliability provides a strong tool to distinguish between the Bayesian model and heuristic-based models in model comparison. Indeed, we found that we were unable to distinguish between the Bayesian and Max models in conditions with fixed stimulus reliability, while the Max model was convincingly rejected in conditions with mixed reliabilities. These results strongly suggest that humans – just like Bayesians – take into account stimulus reliability during perceptual decision-making. This finding is consistent with previous studies that have drawn a similar conclusion in the context of not only visual search [13], but also categorization [18], change detection [19], and same/different discrimination [21] tasks. However, unlike those previous studies, we do not interpret this finding as evidence for near-optimality, because we also found evidence for substantial suboptimalities that are seemingly caused by computational imperfections.

### Suboptimality in perceptual decision making

Although reports of optimality have dominated perceptual decision-making literature, we are certainly not the first to report evidence for suboptimalities. For example, numerous sensory cue combination studies have reported overweighting of one of the sensory cues [50–57]; Bhardwaj et al. [58] found that visual search performance is suboptimal when stimuli are correlated; Ackermann and Landy [59] reported that subjects fail to maximize reward in a visual search task with unequal rewards across target locations; and Qamar et al. [60] found that both humans and monkeys performed suboptimally in a relatively simple visual categorization task. However, none of those studies used the factorial modeling design that we proposed and, therefore, could not distinguish between suboptimalities due to a fundamentally wrong decision strategy and suboptimalities due to computational precisions.

### Late noise in models of perceptual decision making

An important aspect of our analysis is that we included models with “late noise” on the decision variable. We are not the first to do so. An example of our own previous work – in which we referred to it as “decision noise” – is the change detection study by Keshvari et al. [19], where we found that inclusion of late noise did not substantially improve the model fits. However, sensory noise levels in that study were fitted in an entirely unconstrained way, while it is conceivable that there was a trade-off between effects of noise on the decision variable and effects of sensory noise on model predictions. Moreover, in that study we assumed random variability in encoding precision, which a later study showed may be confounded with decision noise [61]. Therefore, it is possible that computational imperfections in the study by Keshvari et al. went unnoticed due to confounding them with sensory noise or variability in precision.

Another body of work that has considered noise on the decision variable are the studies by Summerfield and colleagues (e.g., [62,63]). They have shown that in the presence of late noise, subjects can – and often do – obtain performance benefits by using “robust averaging”, i.e., down-weighting outlier cues when computing the global decision variable. From an optimal-observer perspective, our task can also be conceived of as an averaging task, even though the averaging is over local posterior evidence values, Eq. (2), rather than directly over stimulus values. We performed simulations to examine whether robust averaging also gives performance benefits in our task, but we did not find any evidence for this.

While late noise seems to be an important factor in explaining behavior on our visual search task, it seems to play no role in explaining behavior on classical cue combination tasks [12,56]. There are two differences between these tasks that may explain the difference in findings. First, subjects in our task had to combine four cues instead of two. Second, and perhaps more importantly, the optimal decision rule in our task is substantially more complex: while optimality on cue combination tasks can be achieved using only linear operations, our visual search task required non-linear computations, Eq. (2). Previous work has suggested that information processing in the human brain proceeds mostly by linear additive integration (e.g., [64,65]), which would lead to suboptimalities if the optimal strategy requires non-linear computations. It would be interesting to investigate in future work whether subjects are perhaps using linear approximations to optimal decision rules in complex tasks such as visual search.

### The effect of external uncertainty on performance

A difference between most laboratory stimuli and naturalistic stimuli is that the former are typically deterministic, while the latter are often probabilistic [32]. In the present study, we mimicked the probabilistic character of naturalistic stimuli by adding external uncertainty. While we are not the first to do so in a perceptual decision-making task (e.g., [27,60,66,67]), we are unaware of any previous work that has systematically varied this level of uncertainty to them. Moreover, previous work did not examine the relation between the magnitude of the external uncertainty and the magnitude of deviations from optimality. None of our analyses provided evidence that external uncertainty affects how much performance deviates from optimality. This is somewhat surprising, because the stimulus distributions in our experiment were arbitrary and entirely novel to our subjects. A possible explanation could be that the brain may be familiar with Gaussian-like stimulus ambiguity and can therefore quickly incorporate novel kinds of external uncertainty, as long as it follows a Gaussian distribution. An interesting direction for future work would be to further investigate the relation between different types of external uncertainty and optimality in human decision making.

## S1 Appendix: Supplementary Methods

### Derivation of the optimal decision variable for the visual search task

We denote target presence by a binary variable *T* (0=absent, 1=present), set size by *N*, the mean of the target distribution by μ_T_, the width of the stimulus distributions by *σ*_ext_, the stimulus values on a given trial by **s**={*s*_1_, *s*_2_, …, *s*_*N*_}, the location (index) of the target by *L*, and the observer’s noisy observations of the stimulus values by **x**={*x*_1_, *x*_2_, …, *x*_*N*_}. The Bayesian optimal observer reports “target present” if the posterior probability of target presence exceeds that of target absence, *p*(*T*=1|**x**) > *p*(*T*=0|**x**). This strategy is equivalent to reporting “target present” if the log posterior ratio exceeds 0,

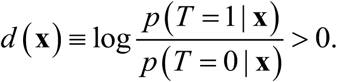

where *d*(**x**) is referred to as the decision variable. Applying Bayes’ rule, we find

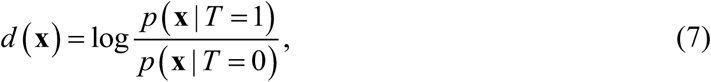

where we made use of the fact that *p*(*T*=0)=*p*(*T*=1) in all our experiments. Using basic rules of probability, we rewrite the numerator to

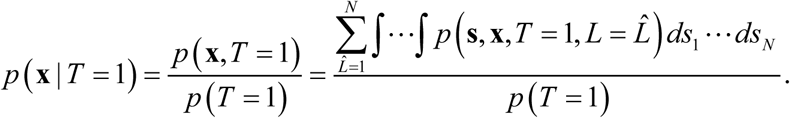

Applying our knowledge of the generative model (S1 Fig), this evaluates to

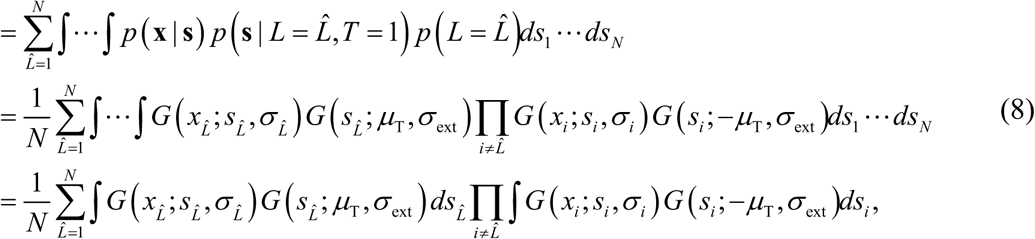

where *G*(*x*; μ, *σ*) is a Gaussian distribution with mean μ and standard deviation *σ*. The denominator of Eq. (7) evaluates to

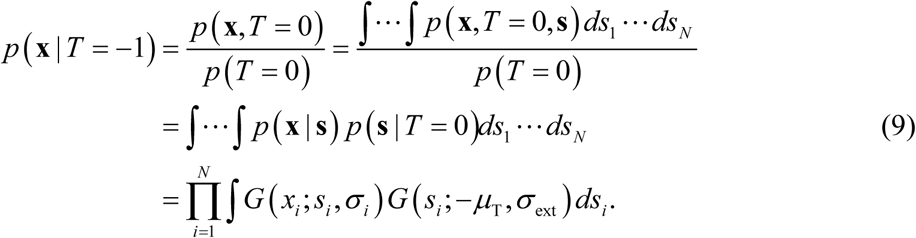

Combining Eqs. (7), (8), and (9) yields the decision variable as presented in the main text:

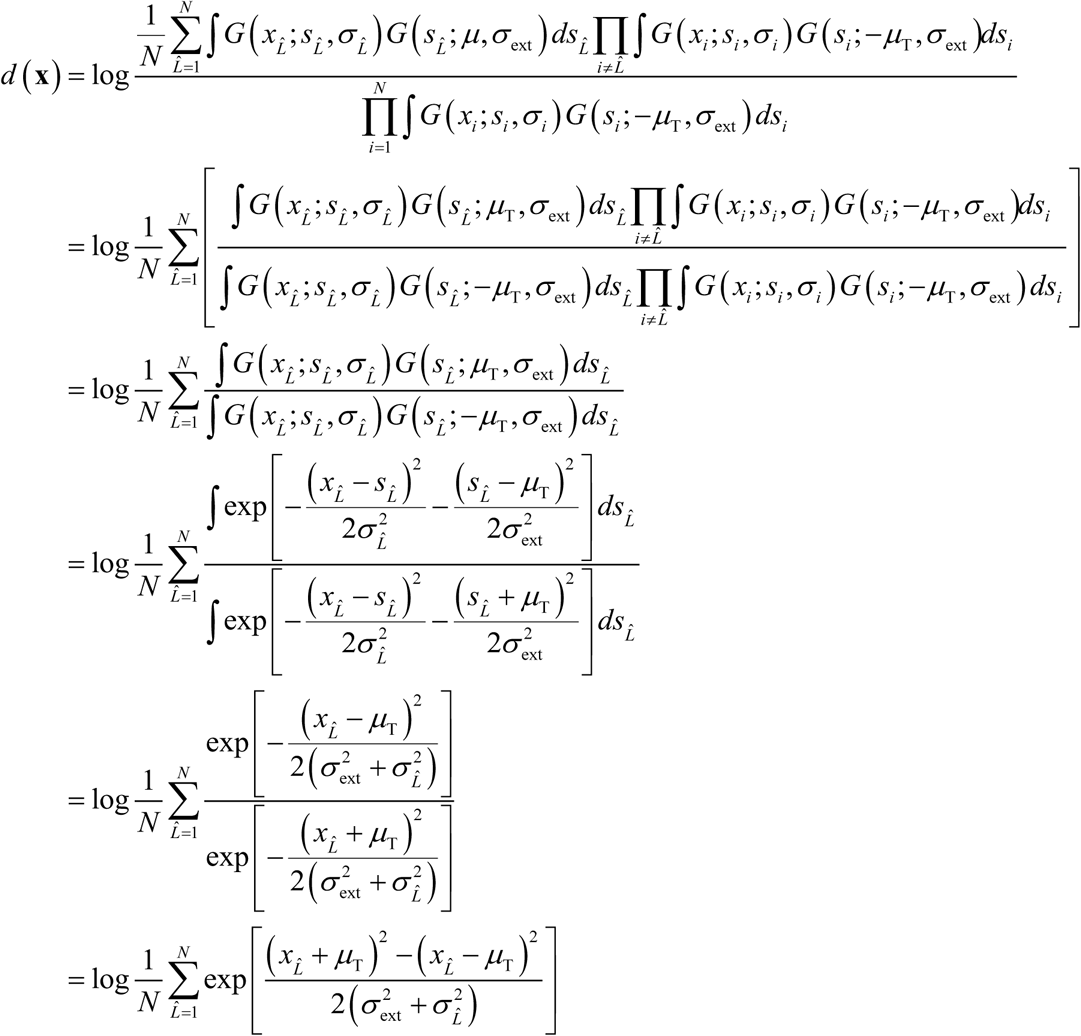

### Constrained fitting of the lapse rate parameter

If we do not constrain the lapse rate parameter in the Bayesian models, then these models may explain away decision suboptimalities by overestimating lapse rates. To avoid this unwanted flexibility, we use data from the discrimination task to obtain independent estimates of subjects’ lapse rates and then use this information to constrain lapse rates in the models for the visual search task. The logic behind this approach is that lapse rates in the discrimination task can be estimated quite accurately and there is no reason to assume that the frequency of attentional lapses is very different in the visual search task. (One may argue that subjects guess more often when the task is more difficult. However, a difficulty-driven increase in guessing is likely to be due to a higher frequency of non-informative decision variables, rather than due to a higher frequency of attentional lapses. Models should account for this by processes that deteriorate the decision variable, not by increasing the frequency of attentional lapses.) We estimate each subject’s lapse rate by fitting a cumulative Gaussian with a free lapse rate parameter to their discrimination task data. According to these estimates, subjects guessed on 3.2±1.2% of the trials. We fit a beta distribution to the distribution of all individual lapse rate estimates and use this distribution as a prior on the lapse rate in the Bayesian models for the visual search task. The parameters of the beta distribution are α=0.18 and *β*=5.61.

### Constrained fitting of sensory noise level parameters

If we fit sensory noise parameters *σ*_low_ and *σ*_high_ in an entirely unconstrained manner, then we may be giving models an opportunity to explain away decision suboptimalities by overestimating sensory noise levels. We can reduce such flexibility by constraining parameters *σ*_low_ and *σ*_high_ with prior information about the expected sensory noise levels in our experiment. Prior to performing the visual search experiment, each subject performed a discrimination task with an ellipse stimulus identical to the ones used in the visual search task. Due to the simplicity of the discrimination task, our estimates of *σ*_low_ and *σ*_high_ in that task (Table 2 in main text) are probably highly representative of the true noise levels in that task. Unfortunately, however, these estimates may not be representative for the noise levels in the visual search task, because it used a different set size (4 instead of 1) and previous work has shown that sensory noise levels may increase with set size, up to a factor of two [1,2]. We express this increase as ratios, 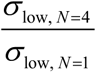 and 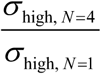. To get an estimate of a plausible range for these ratios, we performed a control experiment that includes discrimination tasks at both set sizes 1 and 4.

#### Control experiment

Twelve subjects performed the original discrimination task (as described in the Main text) and a variant of the task with four stimuli. In the variant, subjects were presented on each trial with four stimuli with mixed reliabilities, such that the visual characteristics were identical to the visual search task. One of the stimuli would disappear after 67ms and the task of the subject was to indicate the direction of tilt of the disappeared stimulus. Hence, just as in the visual search experiment, subjects had to encode four stimuli on each trial, because they would not know which of them would disappear. Each subject was tested on two versions with set size 4. In the first version, the stimuli were representative of the visual search task with 0% uncertainty (“lowest heterogeneity”). In the second version, stimuli were representative of the visual search task with 15% uncertainty (“highest heterogeneity”). Each subject performed 300 trials of the original discrimination task and 300 trials of each version of the variant with a set size of 4. The probed stimulus had low reliability in half of the trials and high reliability in the other half.

#### Constructing the prior distributions on σ_low_ and σ_high_

We fitted cumulative Gaussians to each subject’s data to obtain estimates of *σ*_low_ and *σ*_high_ in all three versions of the discrimination task. Thereafter, we computed for each subject the ratio between the estimated noise levels at set sizes 4 and 1. Across all estimates, we find that the average ratio was 1.36 with a standard deviation of 0.39. A Bayesian ANOVA provided no strong evidence for an effect of reliability (low *vs.* high; BF_inclusion_=0.80) or heterogeneity (lowest *vs*. highest; BF_inclusion_=1.36) on the estimated ratio. Therefore, we will impose a single prior distribution to all sensory noise parameter estimates, regardless of stimulus reliability or level of external uncertainty. We choose to do this by using a Gaussian prior with a mean of 1.36 and a standard deviation of 0.39 on the ratio between the noise levels in the visual search models and the estimated noise levels from the discrimination task.

## SUPPLEMENTARY FIGURES

**S1 Figure.**
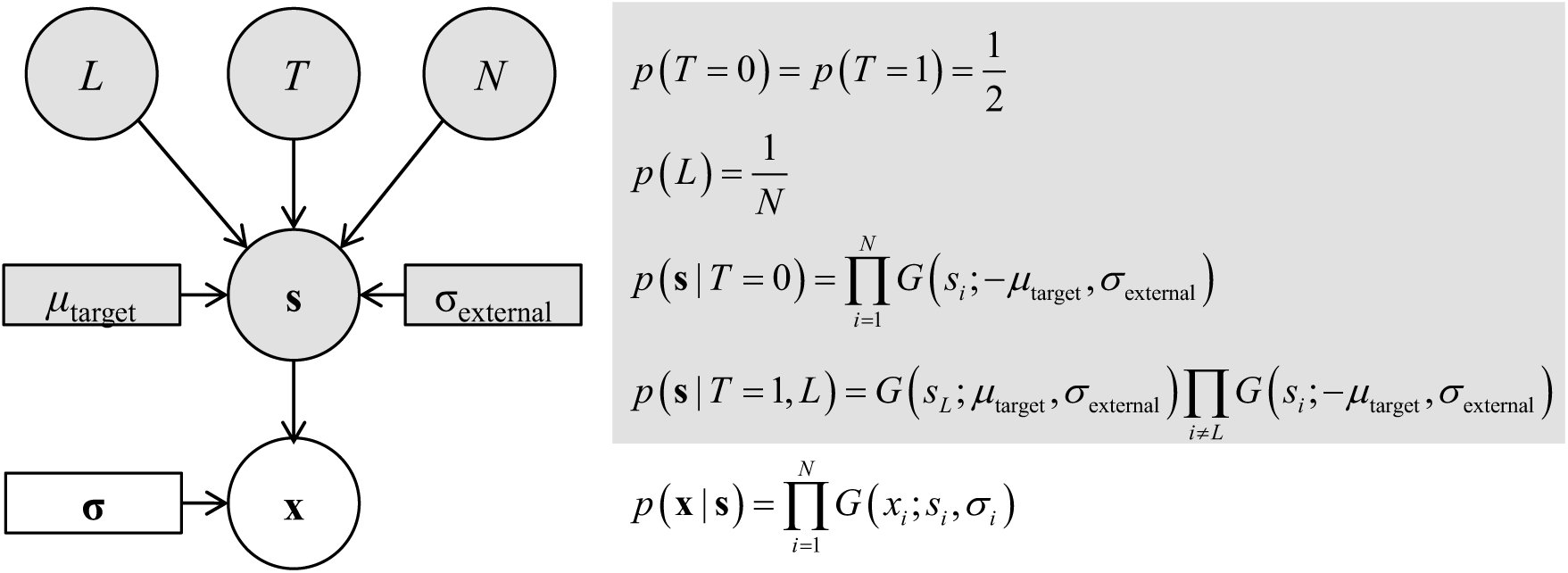
Generative model for the visual search task. Circles represent random variables, rectangles represent constants, and arrows represent causal relationships. Gray shades represent variables and constants that are under control of the experimenter. On each trial, *N*=4 stimuli are presented to the observer. A target is either absent (*T*=0) or present (*T*=1) among these stimuli. Each location *L* ϵ {1, 2, …, *N*} has equal probability of containing the target on target-present trials. On target-absent trials, each stimulus orientation, *s*_*i*_, is drawn from the distractor distribution, which is a Gaussian with a mean −μ_target_ and a standard deviation *σ*_external_. On target-present trials, the stimulus at the target location is drawn from a Gaussian distribution with mean μ_target_ and standard deviation *σ*_external_, while the remaining *N−*1 stimuli are drawn from the distractor distribution. We assume that stimulus observations are corrupted by Gaussian noise, such that each observation, *x*_*i*_, is a Gaussian random variable with mean *s*_*i*_ and standard deviation *σ*_*i*_.

**S2 Figure.**
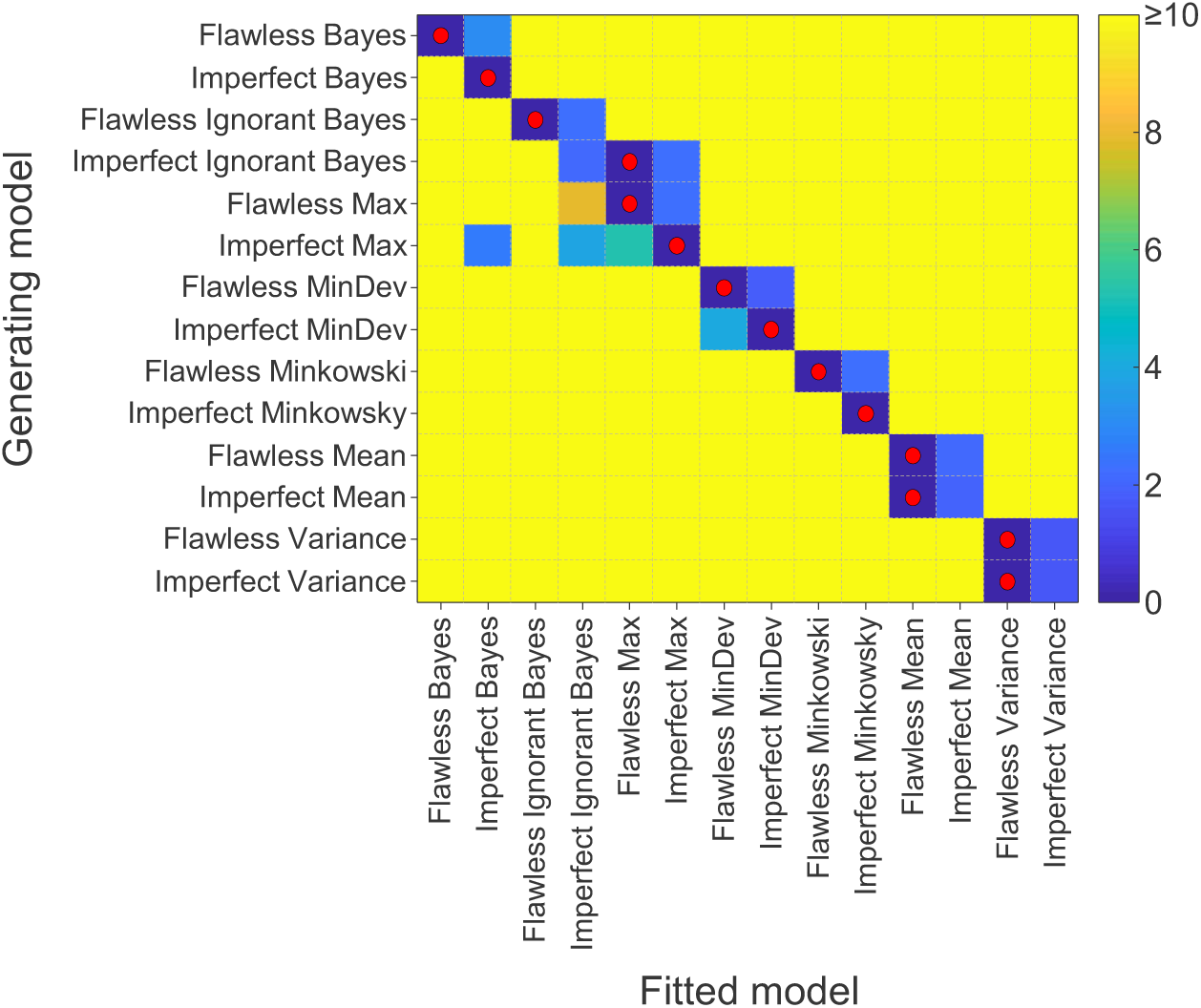
Model recovery results. Ten synthetic datasets were generated from each model, by simulating its responses in trials from the condition with 10% external uncertainty. Each dataset had the same number of trials as a subject dataset. Parameter values were drawn from a multivariate Gaussian distribution with the same mean and covariance as the maximum-likelihood estimates obtained from fitting subject data. Hence, the synthetic datasets had the same size and similar statistics as empirical datasets. Each model was fitted to each of the 140 synthetic datasets. The matrix shows for each generating model the average AIC value (across all ten generated datasets from the model) relative to the best-fitting model. In each row, the overall best-fitting model is indicated with a red dot. In most cases, the generating model is the best-fitting model (red dots on diagonal) and most other models are rejected. There are a few wrong classifications (red dots off-diagonal), which indicates that some model pairs cannot reliably be distinguished from each other. Importantly, the model that was most successful in accounting for subject data – the Imperfect Bayesian model – never is selected as the preferred model when data were generated from another model. Hence, it is unlikely that the success of the Imperfect Bayesian model on empirical data was caused by it being overly flexible.

**S3 Figure.**
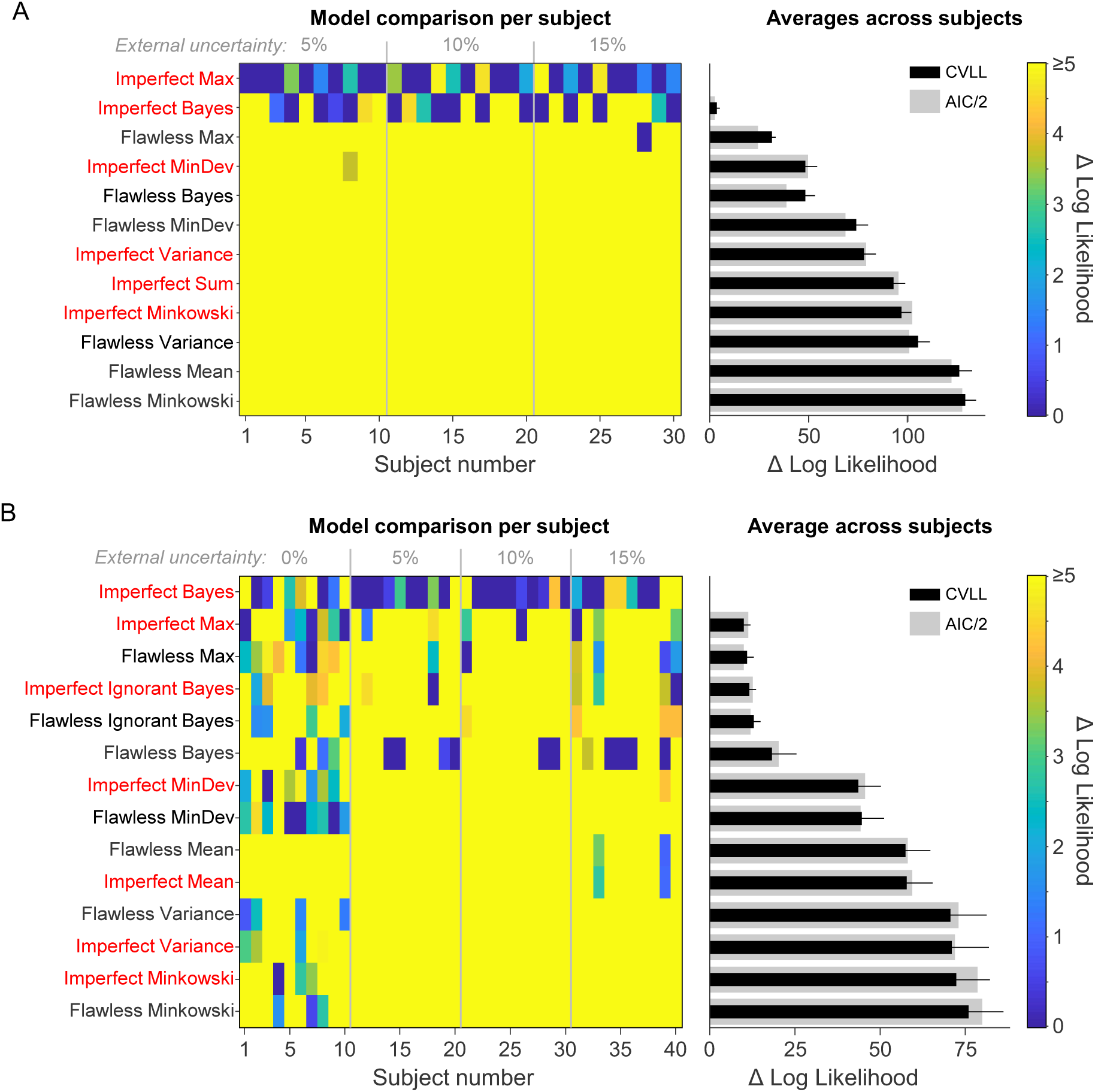
Model comparison results based on five-fold cross validation. Each individual dataset was fitted 5 times. In each of these fits, a different subset of 20% of the trials was left out. The log likelihood of these left out data were computed using the maximum-likelihood estimates obtained from fitting the other 80% of the data. We summed the 5 log likelihood obtained for each subject to compute a single “cross-validated log likelihood” (CVLL). (A) Results from fitting the conditions with unlimited display time. (B) Results from fitting the conditions with brief display time.

